## Supplementary Material for "Cortical beta oscillations map to shared brain networks modulated by dopamine"

| Lobe | Channels per lobe | Sum Theta Channels | Sum Alpha Channels | Sum Beta Channels | Theta Total Peaks | Alpha Total Peaks | Beta Total Peaks |
|---|---|---|---|---|---|---|---|
| **Frontal** | 494 | 48 \| 9.7% | 31 \| 6.3% | 381 \| 77.1% | 190 | 108 | 679 |
| **Temporal** | 451 | 86 \| 19.1% | 158 \| 35.0% | 192 \| 42.6% | 114 | 227 | 668 |
| **Cingulate** | 116 | 12 \| 10.3% | 21 \| 18.1% | 71 \| 61.2 % | 28 | 48 | 150 |
| **Occipital** | 94 | 13 \| 13.8% | 53 \| 56.4% | 27 \| 28.7% | 20 | 59 | 150 |
| **Amygdala** | 3 | 1 \| 33.3% | 0 | 1 \| 33.3% | 2 | 0 | 3 |
| **Insula** | 126 | 18 \| 14.3% | 21 \| 16.7% | 85 \| 67.5% | 29 | 39 | 197 |
| **Parietal** | 220 | 33 \| 15.0% | 65 \| 30.0% | 113 \| 51.4% | 57 | 103 | 369 |
| **Subcortex** | 10 | 1 \| 10.0% | 1 \| 10.0% | 7 \| 70% | 4 | 2 | 16 |

**Supplementary table 1 | sEEG channels** (high and low beta calculated separately to calculate total beta peaks)

| Lobe | Channels per lobe | Sum Theta Channels | Sum Alpha Channels | Sum Beta Channels | Theta Total Peaks | Alpha Total Peaks | Beta Total Peaks |
|---|---|---|---|---|---|---|---|
| **Frontal** | 122 | 25 \| 20.5% | 7 \| 5.7% | 77 \| 63.1% | 48 | 38 | 162 |
| **Temporal** | 68 | 20 \| 29.4% | 22 \| 32.4% | 24 \| 35.3% | 20 | 36 | 102 |
| **Occipital** | 13 | 2 \| 15.4% | 9 \| 69.2% | 0 | 2 | 9 | 19 |
| **Insula** | 2 | 1 \| 50% | 0 | 1 \| 50% | 1 | 0 | 3 |
| **Parietal** | 52 | 17 \| 33.0% | 8 \| 15.4% | 26 \| 50% | 19 | 16 | 80 |
| **Subcortex** | 1 | 0 | 1 \| 100% | 0 | 0 | 1 | 1 |

**Supplementary table 2 | ECoG channels. Lobes with only zero values removed** high and low beta calculated separately to calculate total beta peaks)

### Supplementary Figure 1

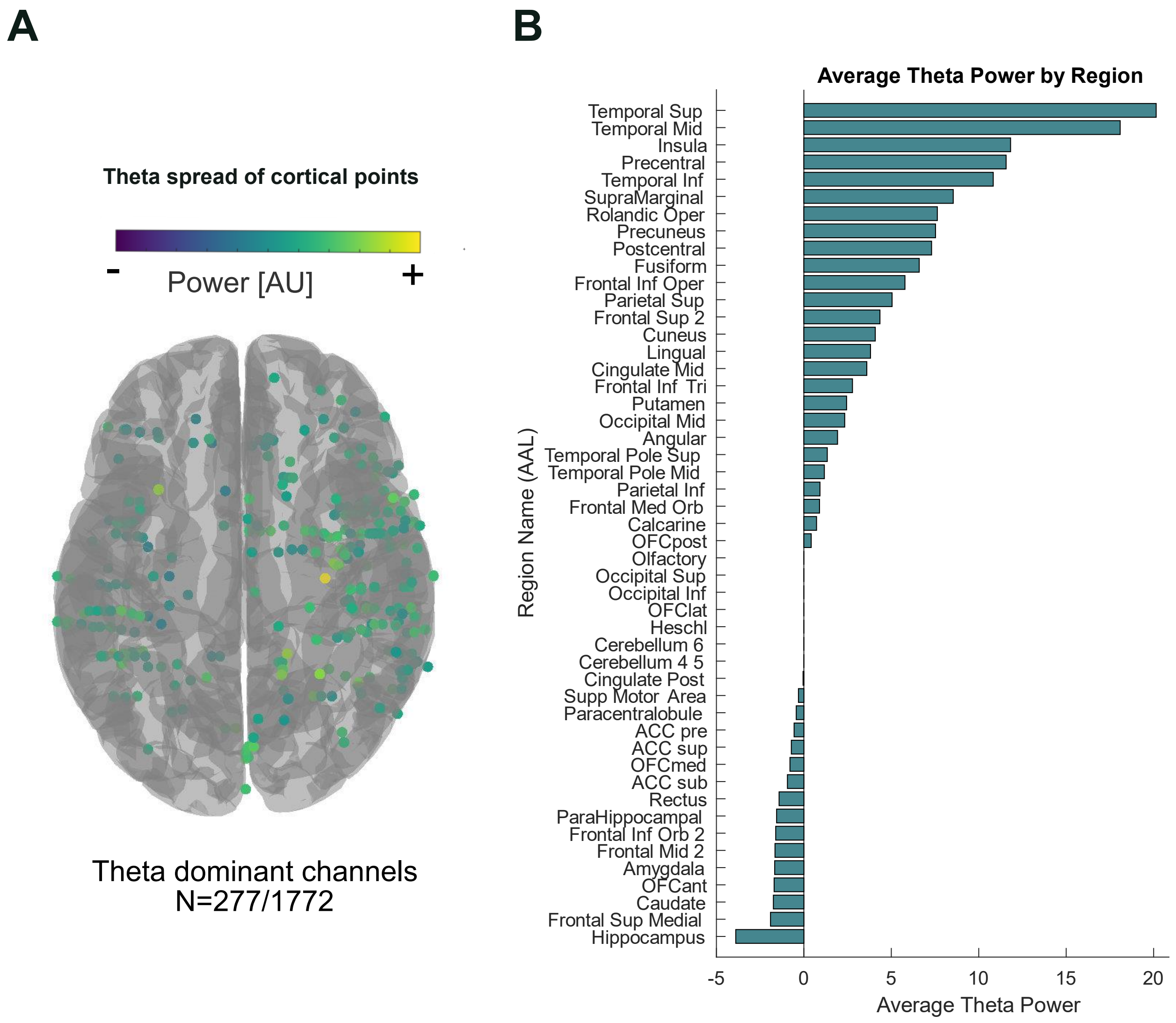


**Figure 1. Cortical spread and amplitudes of theta peaks. (A)** 277/1772 channels showed higher maximum peak power in theta (4 – 8 Hz) than beta (13 – 30 Hz), alpha (8 – 12 Hz) and gamma (30 – 100 Hz). **(B)** Theta dominant peaks were concentrated around the temporal regions however interestingly not in hippocampus. We suspect this is due to the sparse nature of electrodes placed in the region.

#### Supplementary Figure 2

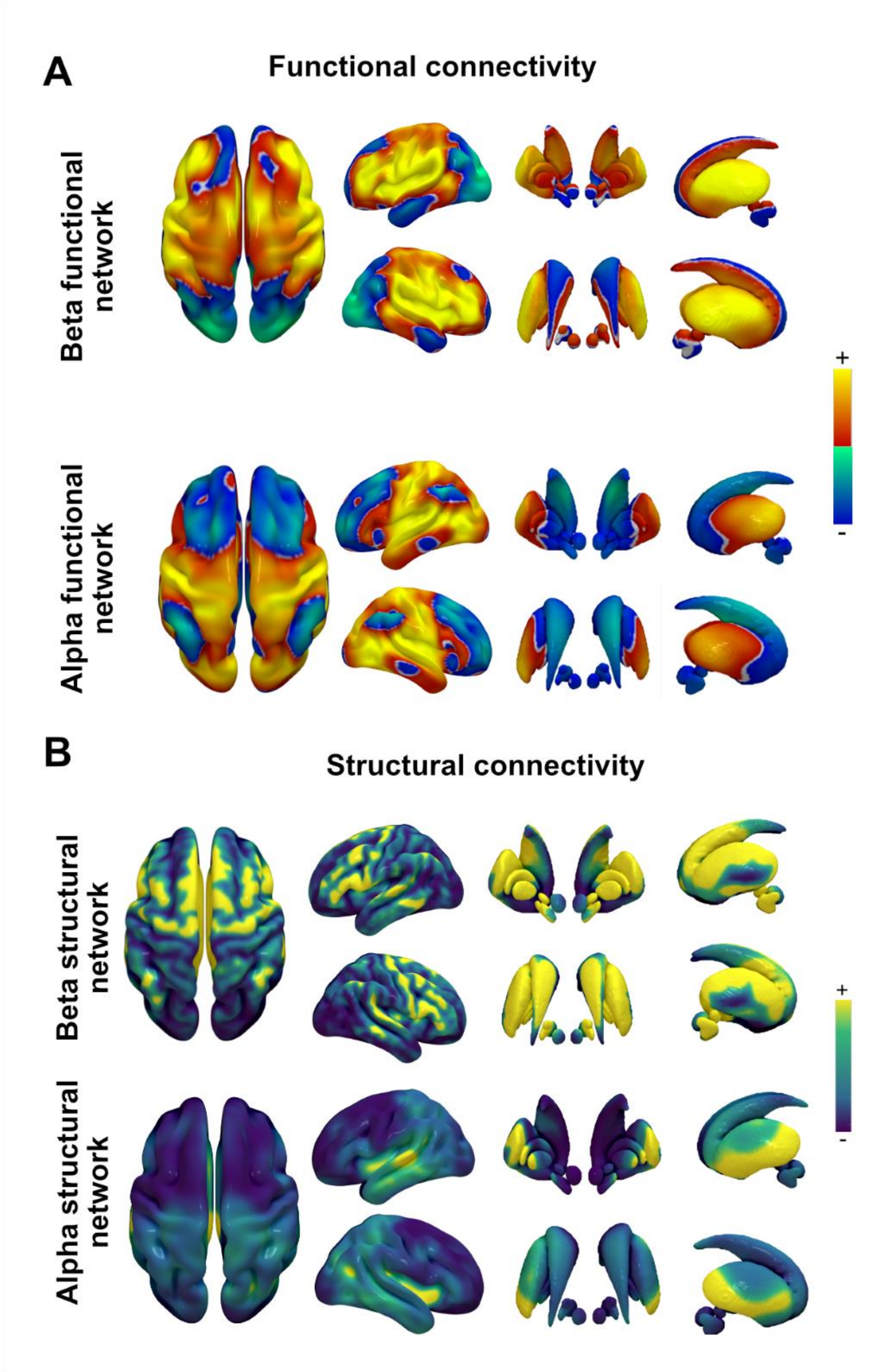


**Figure 2**. **Unthresholded averages of functional and structural connectivity for alpha and beta dominant recording locations**. Functional **(A)** and structural **(B)** group averages without statistical comparison replicate the statistical differences, but show more relative overlap in some regions, e.g. see higher alpha band activity in posterior basal ganglia nuclei.

### Supplementary Figure 3

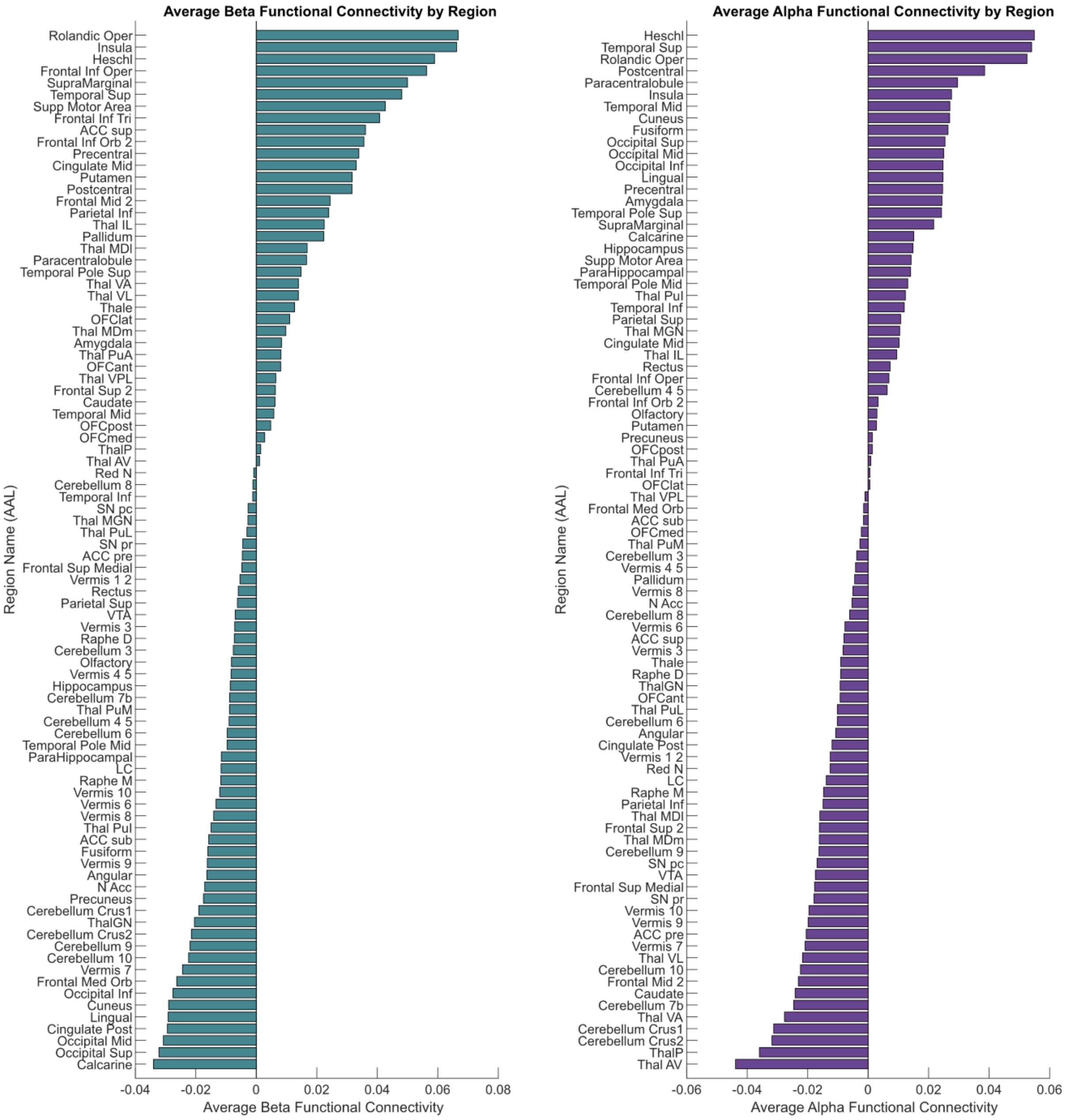


**Figure 3. Region-wise bar plots of functional connectivity values for every parcel based on the automatic anatomical labeling (AAL) atlas**. For functional connectivity, the highest intensities for beta (teal) channels were in frontal, insula, cingulate, basal ganglia and temporal regions, while for alpha (purple) the highest intensities were found in temporal and occipital regions.

### Supplementary Figure 4

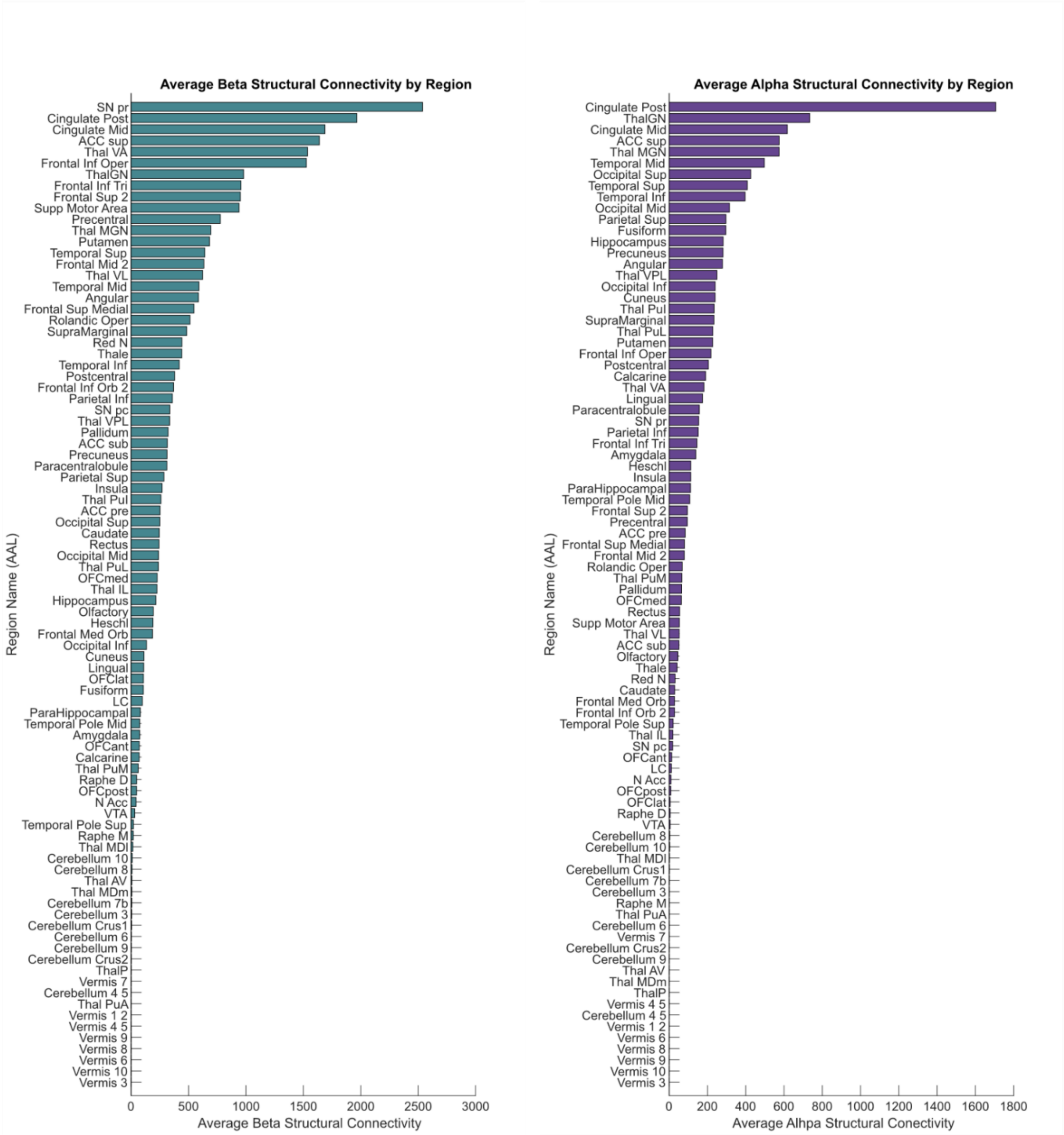


**Figure 4. Region-wise bar plots of structural connectivity values for every parcel based on the automatic anatomical labeling (AAL) atlas**. For structural connectivity, the highest intensities for beta (teal) channels were in basal ganglia, cingulate, frontal, basal ganglia and parietal regions, while for alpha (purple) the highest connectivity estimates were found in cingulate, temporal and occipital regions.

### Supplementary Figure 5

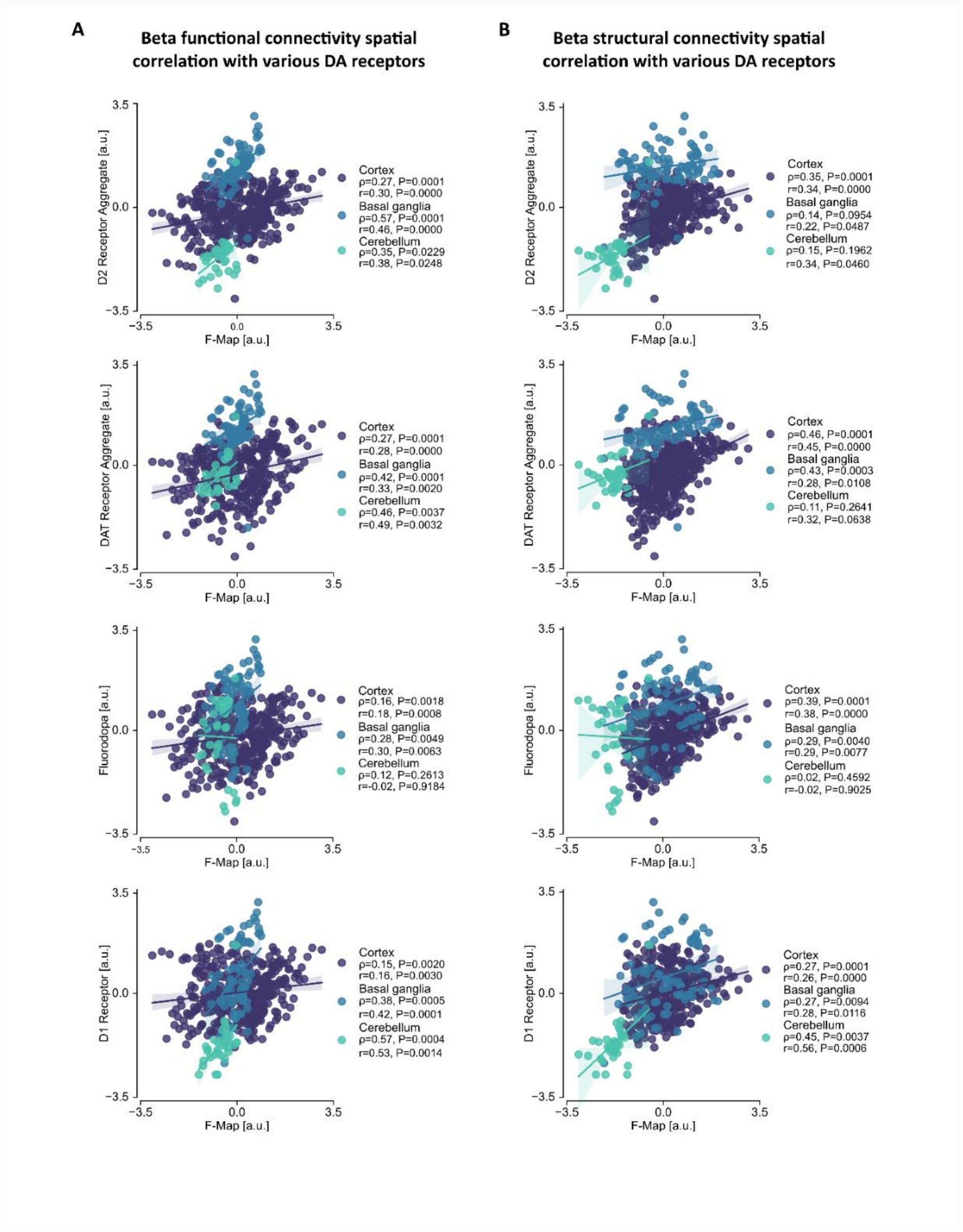


**Figure 5. Correlation plots for individual Dopamine receptors**. **(A)** Beta functional connectivity correlates to an aggregate of the Dopamine receptors as shown in the main article, but also correlates to individual receptors inlcuding D2, the Dopamine transporter protein (DAT), Fluorodopa (FDOPA) and D1. **(B)** Beta structural connectivity for individual Dopamine receptors is significant for each one including D2, DAT, FDOPA and D1.

#### Supplementary Methods

The methods for the invasive electrophysiology atlas1 are as described in the original work.

**Selection of Intracranial EEG Recordings:**

For the inclusion of patients and brain signals in the neurophysiological atlas, archival clinical datasets were reviewed from hospitals across Canada including Montreal Neurological Institute and Hospital, Centre Hospitalier de l'Université de Montréal, and Grenoble-Alpes University Hospital. Selected patients had to have undergone surgical intracranial EEG for epilepsy assessment between January 2010 and April 2016. The following inclusion criteria were used:

**Inclusion Criteria:**

1. **Normal Activity Channel:** The patient had to have at least one channel displaying normal activity as per previous standards used by the group2. The channel had to be in normal tissue, outside of the seizure onset zone and not show any epileptic discharges or slow wave anomalies.
2. **Peri-Implantation Imaging:** Either a CT or MRI scan of post implantation had to be available in order to confirm the localisation of electrodes. All white matter contacts were excluded.
3. **Controlled EEG Recording:** A controlled intracranial EEG was recorded post-implantation was available and adhered to previously used timeframes post-seizure events2.
4. **Sampling Frequency:** The minimum sampling frequency should have been of 200 Hz, aligning with classical Berger frequency bands (0.3–70 Hz).

**Co-registration and Anatomical Localisation of Electrodes:**

Using minctools http://www.bic.mni.mcgill.ca/ServicesSoftware/ServicesSoftwareMincToolKit) and the IBIS framework (https://ibis-project.org/), electrode positions were visualised in a standardised atlas. Peri-implantation CT/MRI images were linearly registered with pre-implantation MRIs, then matched to the ICBM152 2009c non-linear symmetric brain model. Accurate anatomical segmentations were achieved using an atlas of 132 grey matter labels. Each bipolar channel's location was determined based on the presence of the maximum volume present anatomically.

**EEG Sections Selection:**

60-second EEG sections during resting wakefulness that were part of a controlled assessment across centres were chosen for analysis. Signals were filtered (0.5–80 Hz) and appropriately downsampled based on original rates with Matlab.
